# The *Cryptococcus* extracellular vesicle transcriptome

**DOI:** 10.1101/2023.12.08.570810

**Authors:** Adam Taheraly, Corinne Maufrais, Frédérique Moyrand, Jessie Colin, Jean-Yves Coppee, Guilhem Janbon

## Abstract

Extracellular vesicles (EVs) are now recognized as key players in the biology of numerous organisms, including pathogenic fungi. However, studying EVs in these organisms remains challenging. The recent implementation of new protocols to purify EVs in the pathogenic yeast *Cryptococcus neoformans* has resulted in a more detailed description of their structure and protein composition. Although a few publications describing RNA molecules associated with EVs have already been published, we reasoned that these new protocols would be beneficial for gaining a deeper understanding of the EV transcriptome. We thus purified EVs and confirmed that some RNAs were associated with these EV extracts. Iodixanol gradient analyses also revealed that these RNAs co-sedimented with EVs. We then sequenced these RNAs in parallel with RNAs extracted from the very cells producing these EVs using different types of sequencing libraries. Our data confirm the presence of siRNAs and tRFs associated with EVs, some of which are enriched. We also identified some snoRNAs, which in *Cryptococcu*s are mostly borne by coding gene or lncRNA introns.

---

Extracellular vesicles (EVs) are membrane bounded particles released by cells into the extracellular space. EVs have been described in all organisms in which they have been looked for (Van Niel *et al*. 2018). In mammals, in which they have been extensively studied, they have been shown to participate in cell-to-cell communication during development and normal physiology (Van Niel *et al*. 2018). They can also play a positive or negative role in pathological situations such as cancer, diabetes, etc. (Van Niel *et al*. 2022). In most cases, RNAs (mRNAs, tRNA halves, miRNAs, siRNAs…) have been identified as the active ‘message-bearing’ molecules within EVs (Valadi *et al*. 2007). EV RNAs have also been shown to play major roles in host-pathogen interactions (Munhoz Da Rocha *et al*. 2020). Seminal examples in parasitic worms, pathogenic bacteria and plant and insect fungal pathogens have been published in recent years, revealing a widespread usage of this type of interaction during infections (Buck *et al*. 2014; Cui *et al*. 2019; Rutter *et al*. 2022; Sahr *et al*. 2022). Here also, some RNA molecules have been shown to be implicated in these interactions (Weiberg *et al*. 2013; Buck *et al*. 2014; Cui *et al*. 2019; Sahr *et al*. 2022).

EVs have been identified in fungi for the first time in 2007 (Rodrigues *et al*. 2007), and since then they have isolated from every class of fungi (Rizzo *et al*. 2020). In human pathogens, EVs have been shown to play a role in cell intercommunication impacting fungal drug resistance in biofilm or filamentation as well as fungal replication within macrophages and virulence (Bielska *et al*. 2018; Zarnowski *et al*. 2021; Bitencourt *et al*. 2022a; Honorato *et al*. 2022). Yet, the physical bases of the exchanged messages and their mechanism of action remain mostly unknown. By analogy with other systems, one can anticipate that some RNAs could play essential roles. In agreement with this hypothesis, long-distance action of EVs on *Cryptococcus* intracellular replication within macrophages can be abolished by EV RNase treatment (Bielska *et al*. 2018).

*Cryptococcus neoformans* is a pathogenic yeast responsible for 200 000 deaths every year in the world (Rajasingham *et al*. 2017) which has been listed as one of four human fungal pathogens of the “critical group” by the WHO (Fisher and Denning 2023). The production of polysaccharidic capsule, the synthesis of melanin, the production of diverse enzymes associated with its ability to grow at 37°C have been considered as the main *Cryptococcus* virulence factors (Kwon-Chung *et al*. 2014). More recently, EVs produced by pathogenic fungi have been suggested to be also key factors in fungal pathobiology (Rizzo *et al*. 2021a). Nevertheless, EVs studies in *Cryptococcus* have long been limited by cumbersome protocols needed to obtain sufficient EVs even though fungal EVs have been isolated for the first time in this species (Rodrigues *et al*. 2007). Indeed, the initial published protocols used extended cultures (2 to 3 days) in complete liquid medium and often required several liters of culture to obtain enough vesicular extract for a single experiment (Rodrigues *et al*. 2007). This extended duration for cultures also led some to believe that the obtained vesicles actually originated from dead cells with no physiological role (Brown *et al*. 2015). A new protocol published in 2019 by Reis and colleagues in which cells are grown on solid medium for 24h helped resolve this issue and partially address these criticisms (Reis *et al*. 2019). We recently used EVs isolated with this protocol to study the structure of *Cryptococcus* EVs using proteomics, nano flow cytometry, CryoEM/TM and genetic analyses (Rizzo *et al*. 2021b). These experiments revealed a spectacular diversity in size, shape and structure of *Cryptococcus* EVs and helped us to propose a structural model of *Cryptococcus* EVs in which these extracellular organelles are surrounded by capsular polysaccharide and decorated by GPI-anchored and membrane proteins (Rizzo *et al*. 2021b).

While these analyses have greatly improved our understanding of the structure and protein content of EVs produced by *Cryptococcus*, the other molecules associated with these vesicles, particularly their RNAs, remain superficially described (Rizzo *et al*. 2020; Rizzo *et al*. 2021a). In fact, despite a few published studies (Bitencourt *et al*. 2022b), there is no real consensus on the structure of the fungal EVs transcriptome. This is partly due to the lack of consensus on the protocols to be used to prepare vesicular extracts, but also to the diversity of sequencing and sequence analysis strategies used. For instance, there are three publications describing the results of RNA sequencing from *Cryptococcus* EVs (Peres Da Silva *et al*. 2015; Peres Da Silva *et al*. 2018; Liu *et al*. 2020). Even though these results clearly open up possibilities by describing RNAs associated with fungal EVs, neither the EV purification processes, nor the bioinformatics analyses have been standardized. Moreover, no enrichment analysis has been performed and further experimental validation is often missing. Therefore, it is difficult to estimate the share of simple contaminants within the pool of RNA species identified. Thus, although RNAs associated with fungal EVs have been reported in many species (Bitencourt *et al*. 2022b), the fungal EV transcriptome remains poorly described.

In the present study, we used the Reis protocol to purify EVs from three species of *Cryptococcus* and we studied their transcriptome. Furthermore, we implemented an additional purification step based on iodixanol density gradient to show that RNA molecules co-sediment with EVs. Finally, we used several types of RNA sequencing libraries and bioinformatic strategies to compare the cellular and vesicular transcriptomes in *C. neoformans*.

## Material and Methods

### Strains

The strains used in this study were *C. neoformans* KN99α (Nielsen *et al*. 2003), *C. deneoformans* JEC21 (Loftus *et al*. 2005) and *C. deuterogattii* R265 (Ma *et al*. 2009).

### EV sample preparation

EV extracts were obtained as previously described (Rizzo *et al*. 2021b). Briefly, *Cryptococcus* cells were grown for 16-hour in liquid YPD at 30°C and plated onto 24 minimal SD agar medium dishes. The plates were then incubated at 30°C for 24 hours and scratched to collect the cells, which were then gently resuspended in cold 0.01 M PBS (filtered at 0.02 µm). To separate the supernatant from the cells, two successive centrifugation steps were performed at 5,000 *g* and 15,000 *g* for 15 minutes at 4°C. The resulting cell pellets were snap frozen ethanol/dry ice and stored at -80°C. EVs were then pulled down from the 0.45 µm filtered supernatant by ultracentrifugation at 100,000 *g* for 1 hour at 4°C using a SW 40Ti rotor. The resulting pellet was resuspended in 400 µL of filtered 0.01 M PBS. When necessary, EVs were further purified on 40-5% w/v iodixanol gradient as previously described (Reis *et al*. 2021). After 18 h of ultracentrifugation at 100,000 *g* 4°C, fractions of 1 mL were collected by pipetting to which 9 mL of filtered 0.01 M PBS were added prior ultracentrifugation (1h 100,000 *g* at 4°C). Each pellet was then resuspended in 100 µL of filtered 0.01 M PBS and stored at 80°C.

### RNA extraction

Both EV and cellular RNA extractions were carried out using a TRIzol protocol under RNAse-free conditions at 4°C as previously described (Moyrand *et al*. 2008).

### UREA-PAGE

Heat denatured RNA samples were run in TBE-UREA 8M gel for 70 minutes at 120V. The gel was then washed with water and revealed using SYBRGold (Invitrogen) as suggested by the manufacturer.

### RNA sequencing

For the first strategy, total RNA was treated with Cap-Clip Acid Pyrophosphatase (Cellscript). as previously described (Wallace *et al*. 2020). The sequencing libraries were then constructed using the QIAseq® miRNA Library Kit according to the manufacturer protocol. The sequencing reaction per se were performed using Illumina Iseq100. For small RNA libraries, 20-30 nt RNA molecules were UREA-PAGE purified from both EV and cell samples. RNA was precipitated using a DNA Lo Bind tube (Eppendorf) and recovered in 16.5 µL of water. Triphosphate in 5’ were removed using a RNA 5’ polyphosphatase (Euromedex Lucigen) and RNA were purified using 3 volume of absolute isopropanol and 1.8 volume of Agencourt RNAClean XP beads (Beckman). RNA was eluted with 6 µL of water. After ligation of an adaptater (5’rApp/NNNNTGGAATTCTCGGGTGCCAAGG/3ddC/) in 3’, RNA was purified again. An adaptater (5’GUUCAGAGUUCUACAGUCCGACGAUCNNNN3’) in 5’ was added, and RNA was purified as described previously. RT reactions were performed using 5X SuperScript III (Invitrogen) using conditions suggested by the manufacturer. Libraries were finally amplified using Phusion PCR (10 for cell samples and 14 for EVs samples), checked by TapeStation electrophoresis. UREA-PAGE purified cDNA was then sequenced by Illumina NextSeq 500. Finally, strand-specific, paired-end cDNA libraries were prepared from cell or EV total RNA by polyA selection using the TruSeq Stranded mRNA kit (Illumina) according to manufacturer’s instructions. Then, 100 bases were sequenced from both ends using an Illumina HiSeq2500 instrument according to the manufacturer’s instructions (Illumina). Trimmed reads were mapped to the *Cryptococcus* genome using Bowtie2 (Langmead and Salzberg 2012) and Tophat2 (Kim *et al*. 2013) as previously described (Wallace *et al*. 2020).

### Bioinformatics analysis

Data preparation, analysis, and visualization were conducted using BASH and R 4.3, unless stated otherwise. For the conservation plot, genomic sequences were obtained from FungiDB, including 1000 nucleotides before the start codon and after the stop codon. Homologous sequences from *C. deneoformans* JEC21, *C. deuterogattii* R265, *Kwoniella mangroviensis* CBS 8507, *Kwoniella heveanensis* CBS 569, *Tremella mesenterica* DSM 1558, and *Kwoniella bestiolae* CBS 10118 were identified using blast and retrieved in the same manner. Alignment was performed using MUSCLE 5.1 (Edgar 2004), and visualization was done using JalView 2.11.2.7. Conservation plot data were obtained using plotcon (EMBOSS 6.6) (Rice *et al*. 2000) with a window size of 20 nucleotides.

### Proteomic analysis

Proteomic analysis was performed as described previously (Rizzo et al., 2021). However, in order to eliminate the iodixanol remaining in the gradient purified samples, proteins were first TCA precipitated before being resuspended in PBS and trypsin digested.

## Results

### Some RNAs are EV-associated in *Cryptococcus*

Numerous studies have documented the presence of RNA molecules associated with fungal EVs, including in *Cryptococcus* (Peres Da Silva *et al*. 2015; Peres Da Silva *et al*. 2018; Liu *et al*. 2020; Reis *et al*. 2021; Bitencourt *et al*. 2022b). Nevertheless, the methods of growth, EV purification, and RNA sequencing employed in these studies vary significantly. We also know that the growth conditions can have a strong influence on EV production and EV composition in fungi (Reis *et al*. 2021; Amatuzzi *et al*. 2022; Rizzo *et al*. 2023). This suggested that further analyses would be beneficial for gaining a deeper understanding of the fungal EV transcriptome structure. We here isolated total RNAs from EVs purified from cells cultivated on solid minimum medium, following the established procedure (Reis *et al*. 2021; Rizzo *et al*. 2021b). We then loaded these isolated RNAs onto denaturing urea polyacrylamide gel (Urea-PAGE) alongside RNAs obtained from the cells generating these EVs (Figure 1A). As anticipated, SYBR Gold staining confirmed the presence of RNAs in both cell and EV extracts, indicating that certain RNA molecules are associated with EVs in *C. neoformans.* To further validate this association, we deposited the EV extract on an iodixanol density gradient (Ford *et al*. 1994) (Figure 1B). After 18h of ultracentrifugation at 100 000 *g*, we observed a single whitish ring at a centered position within the otherwise transparent gradient (Figure 1B). Examination of twelve gradient fractions using both flow cytometry with a NanoFCM^TM^ apparatus and ergosterol concentration evaluation (Rizzo *et al*. 2021b; Rizzo *et al*. 2023) showed that the majority of EVs are found within the fractions 6, 7 and 8 (Figure 1C). In fact, the 7^th^ fraction, which corresponds to the whitish ring position, exhibited the highest concentration of EVs, strongly suggesting that this ring corresponds to concentrated EVs. We then conducted proteomic analysis on all twelve fractions. The assessment of protein relative abundance using Intensity Based Absolute Quantification (IBAQ) (Schwanhäusser *et al*. 2011) further corroborated this observed pattern (Figure 1D) (Table S1). The previously described most abundant proteins in *C. neoformans* EVs, Mp88 and Vep1, represent here 58% of the total proteins confirming that these proteins are the major EV proteins in *C. neoformans* (Rizzo *et al*. 2021b). We then isolated RNAs from each of the 12 fractions and analyzed these extracts by Urea-PAGE. SYBR Gold staining unveiled that a significant portion of the RNA was linked to fractions 6, 7, and 8, which also contained EVs and EV proteins. Overall, these analyses provide further evidence that the RNAs purified from our initial EV extract are indeed associated with EVs, in line with previous reports (Bitencourt *et al*. 2022b). However, the precise localization of these RNAs, whether inside, on, or at the surface of the EVs, still requires further investigations.

**Figure 1.**
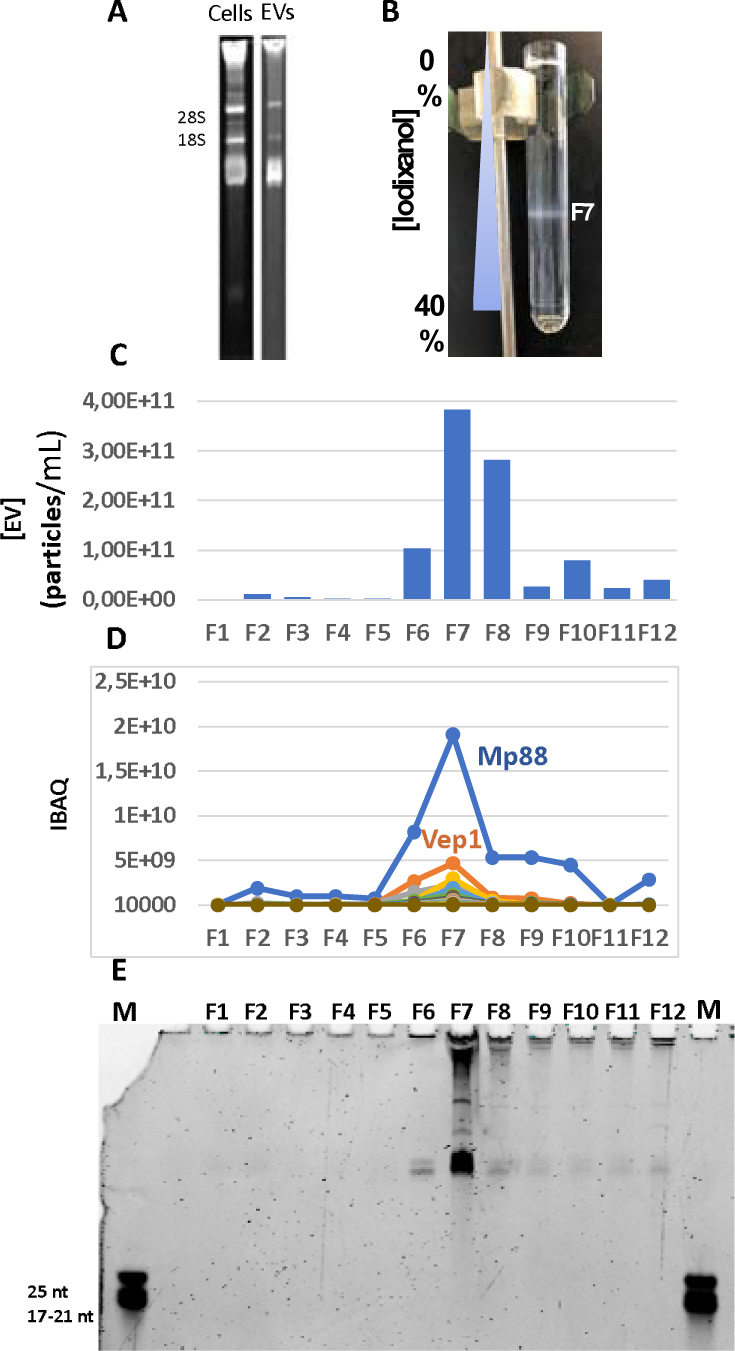
Some RNA molecules are EV associated. (A) Urea-PAGE analysis of cells and EV RNA extracts. (B) Picture of a tube of iodixanol gradient revealing a whitish ring containing EVs around the fraction 7. (C) EV concentration within each fraction evaluated by NanoFCM^TM^. (D) Proteomic analysis of the iodixanol gradient fractions confirming the prevalence of Mp88 and Vep1 in *C. neoformans* EVs. (E) Urea-PAGE analysis of RNAs purified from each fraction of the iodixanol gradient. EV concentrations evaluation and proteomic analyses were performed on the same gradient. RNAs were extracted from samples obtained from an independent experiment.

### Small RNAs associated with EVs

Because a large diversity of RNAs (siRNAs, tRNA halves, lncRNAs, mRNAs, snRNAs, snoRNAs…) have been shown to be associated with EVs in different organisms including fungi (Rizzo *et al*. 2021a; Bitencourt *et al*. 2022b), we first constructed a sequencing library using a small RNA library preparation kit on total RNA extracted from EVs and previously submitted to decapping treatment (Wallace *et al*. 2020), yet no size selection step was implemented at this step. We reasoned that this approach should provide us with a comprehensive view of the *Cryptococcus* EV transcriptome since it should generate small RNA reads as well as, albeit to a lesser extent, longer reads up to 100 nt corresponding to mRNAs or lncRNAs fragments from the same samples. We mapped a total of 1.4 million of reads to the *C. neoformans* H99 reference genome (Janbon *et al*. 2014). A substantial portion of the reads longer than 17 nt (38%) aligned with the ribosomal DNA loci or the mitochondrial genome. The remaining reads exhibited a median size of 26 nt. Among them, the majority (98.3%) mapped to annotated nuclear genome features, either in the sense or antisense orientation. We noticed that most reads aligning in the sense orientation (81.7%) were short reads (median size = 27 nt) aligning to tRNA loci. However, it was obvious that the current annotation of tRNAs was partial with number of them not annotated and several others for which the genomic coordinates were wrong. We thus used tRNAscan (Chan *et al*. 2021) to re-annotate the tRNAs in the *C. neoformans* genome adding 8 pseudo-tRNA loci and updating the coordinates for three tRNAs (CNAG_10023, CNAG_10007, CNAG_10077) (Data S1). The analysis of our sequencing data then revealed that 72.8% of the sense reads mapped to the 3’ and 5’ regions of the annotated tRNAs in *C. neoformans* (Figure 2A, C) suggesting the presence of tRFs (tRNA fragments) in the transcriptome of *C. neoformans* EVs. These tRFs have previously been identified as associated with EVs from different organisms (Lambertz *et al*. 2015; Chiou *et al*. 2018; Artuyants *et al*. 2020; Gámbaro *et al*. 2020). In mammals, tRFs hold promise as biomarkers and have been shown to play diverse regulatory roles in various cancers, neurodegenerative diseases, and responses to infections (Pandey *et al*. 2021). Some tRFs have been shown to associate with Argonaute protein (Ago) in different organisms and several studies reported the role of tRFs in the RNA interference pathway (Maute *et al*. 2013; Ren *et al*. 2019; Sharma *et al*. 2023). Reads aligned to tRNA sequences have already been detected in EV transcriptomes including in human fungal pathogens (Peres Da Silva *et al*. 2015; Munhoz Da Rocha *et al*. 2021). Yet, tRFs functions in these organisms remain to be identified (Jöchl *et al*. 2008). Similarly, although our analysis revealed that a subset of these tRNAs are overrepresented (Pro-tRNA (CGG and UGG), Leu-tRNA (AAG), Gly-tRNA (UCC), Gln-tRNA (CUG) and Asp-tRNA (GUC)) (Figure 2C), the mechanisms underlying this enrichment remain unknown. We performed the same experiment in *C. deneoformans* and *C. deuterogattii* and observed similar pattern at the tRNA loci.

**Figure 2.**
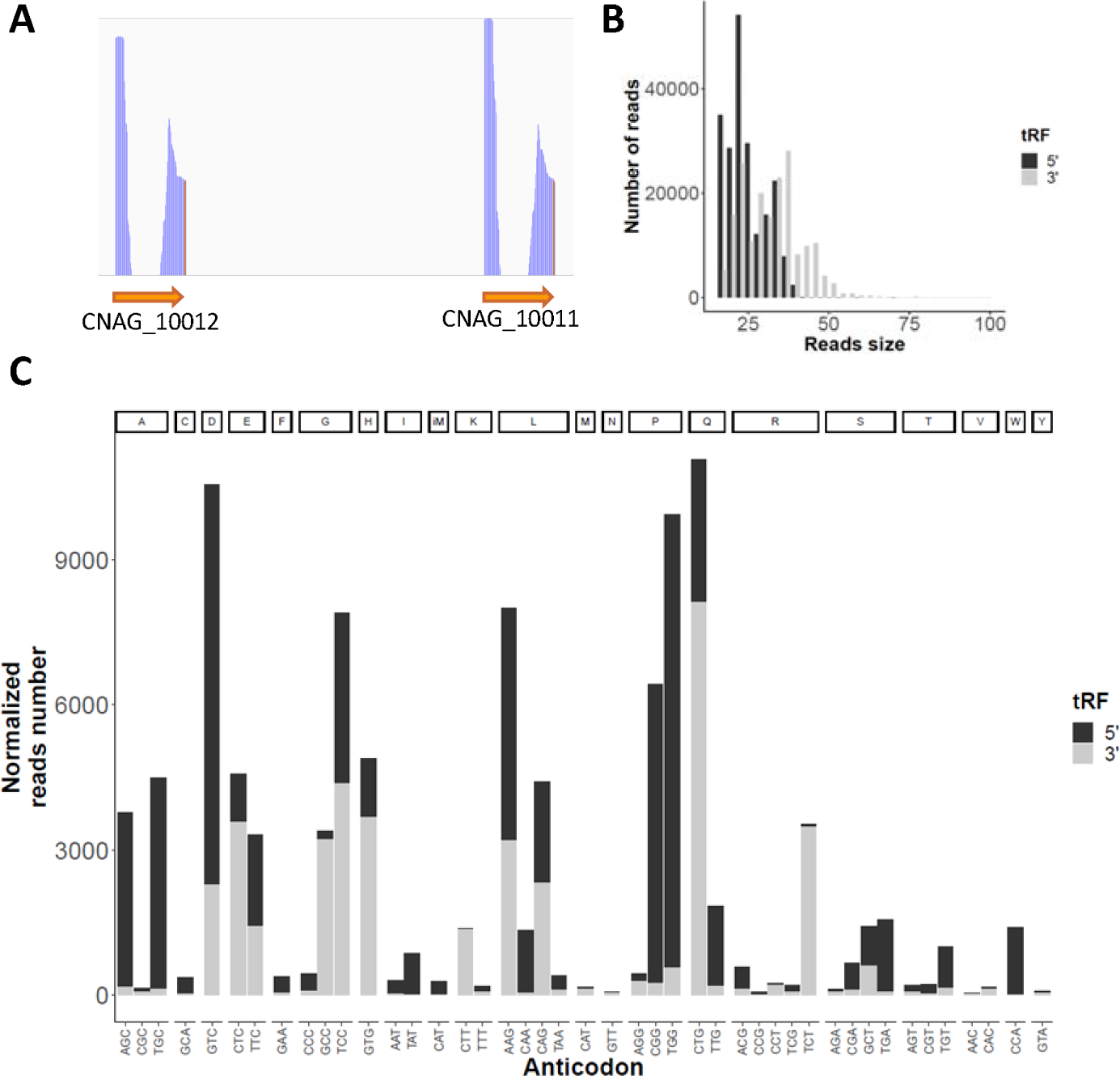
tRFs are associated with *Cryptococcus* EVs. (A) Example of RNA-seq pattern at tRNA loci as visualized by IGV at the loci CNAG_10012 and CNAG_10011 both encoding tRNAs. (B) Number and size of small reads aligned to tRNA loci. (C) RNA-seq reads number associated with the different tRNAs in *C. neoformans*.

By contrast, the examination of reads mapping in the antisense orientation to annotated features showed a significant enrichment of reads ranging from 20 to 25 nucleotides in length, all starting with a uridine. These features are typical of small interfering RNAs (siRNAs) (Figure 3 A, B) (Dumesic *et al*. 2013). As expected, these reads are antisense of transposons or transposon-like sequences (Janbon *et al*. 2010; Dumesic *et al*. 2013) (Table S2).

**Figure 3.**
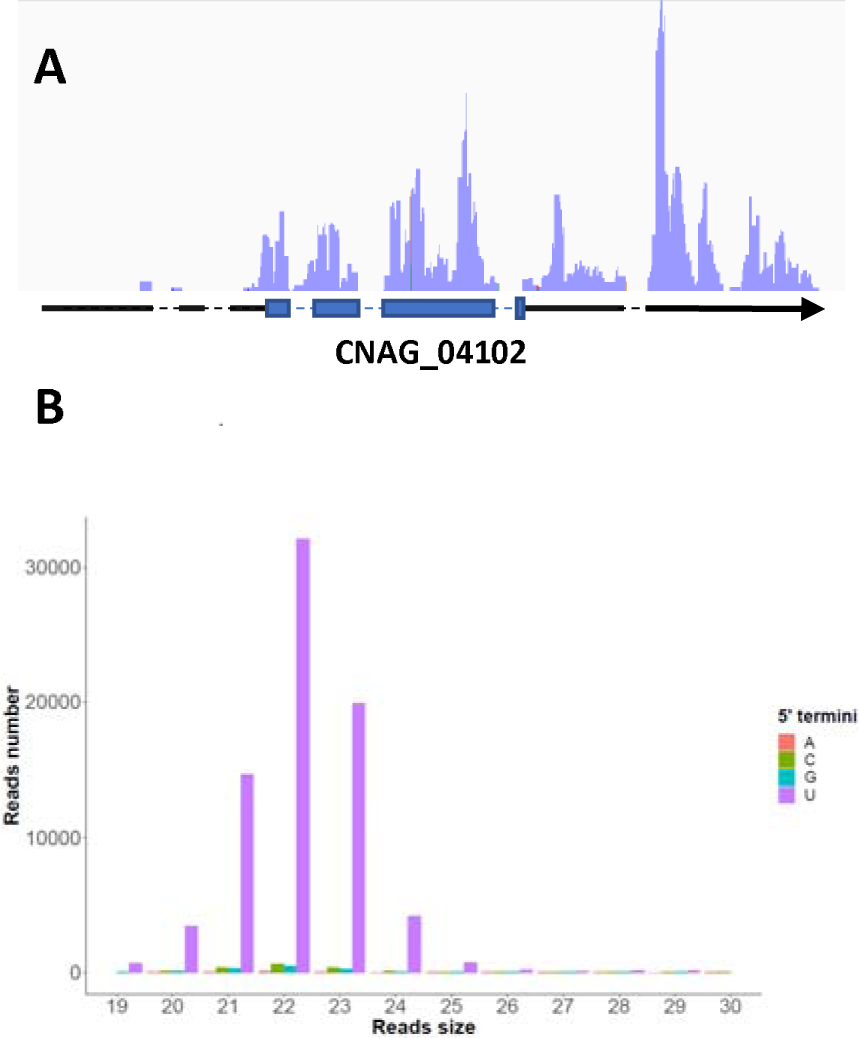
siRNAs are associated with EVs in *C. neoformans*. (A) IGV visualization of antisense reads alignment at the CNAG_04102 locus. (B) Bar plot graph depicting the number of antisense reads as a function of their size and the nature of the 5’ terminal nucleotide.

Additionally, we observed a significant number of these siRNA-like reads mapping within the centromere of all chromosomes, which is consistent with the documented transposon-richness of the centromere structure in *Cryptococcus* (Yadav *et al*. 2018). We performed the same experiment using EVs extracted from *C. deneoformans* and *C. deuterogattii*. The small RNA profile in *C. deneoformans* EVs was very similar to the one obtained in *C. neoformans*. By contrast, as expected, in the RNAi-deficient species *C. deuterogattii*, no siRNA-like species were detected in EVs transcriptome (not shown).

To assess whether the EV small RNA transcriptome differs from that of the cells, we purified on gel 20-30 nt small RNAs from both EVs and cells producing EV to construct small RNA-seq libraries. This experiment was conducted in triplicate and 3.8 to 5.6 million reads obtained from EV libraries and 8 to 9.9 million reads from cell libraries were aligned to the *C. neoformans* genome. Again, most of the reads (98% and 98.6% in EVs and cells, respectively) were either tRFs or siRNAs although their proportion was not identical to the one obtained through sequencing RNAs without implementing a size selection step. By counting reads aligning on arbitrarily defined 500 bp genomic regions, we identified 86 regions containing annotated features with abundant normalized read counts (>100 reads) enriched or depleted in the EV samples compared to the cell samples (Figure 3A) (Table S3). At eight of these loci, the coverage was due to GAU repeat microsatellite-like sequences that were enriched up to 514-fold in EVs. However, the presence of similar reads in the sequencing data, presenting the same repeats but in larger numbers, and which did not align to the genome might suggest that these reads could be artifacts. The most enriched region was specific for reads nearly exclusive of the EVs with an 897-fold enrichment. This 21-nt long read looks like a siRNAs (5’ TTTCACCAACGCCGCTACCAG 3’) and aligned in sense to the Rrp36 ribosomal protein encoding gene at the CNAG_03284 locus (Table S3). Moreover, 74 of these EV enriched regions correspond to tRFs (Table S3). Interestingly, at 36 tRNA loci, the ratio of 5’ tRF vs 3’ tRF reads numbers was statistically different between the cell and the EV samples. For instance, at the CNAG_10094 locus, the 5’tRF is 9.4-fold more abundant in EVs than in the cells whereas the amount of the 3’tRF reads is barely modified (0.8-fold) (Fig 3B). A similar pattern although reverse was observed at the CNAG_10061 locus. Overall, this analysis revealed the presence of siRNAs and tRFs associated with EVs in *Cryptococcus.* The specific enrichment of a subset of small RNAs in EVs suggests a specific mechanism of targeting. In agreement with this model the single Argonaute protein Ago1 was identified in the EV proteome (Table S1)

### SnoRNAs and other RNAs of intermediate size are associated with EVs

The analysis of the sequencing data obtained from total EV RNA samples also identified long reads (>70 nt) (2%, n = 11928) aligning to the *C. neoformans* genome. These reads aligned to 105 precise genomic regions. Most of them (n = 101) correspond to annotated regions, lncRNAs or coding genes, and, surprisingly, a large proportion of them (n = 61) were positioned within introns (Figure 5A). Although these regions did not resemble to any sequence with a known function available in the databases, iterative alignments using BLAST and structure analyses using the RFAM database (Kalvari *et al*. 2021) suggested that number of these sequences are snoRNAs. Further analyses of the sequences data identified 67 snoRNAs with C/D boxes, displaying a classical kink-turn (stem-bulge-stem) structure and characterized by two main terminal consensus sequences (*i.e.* the C box (RUGAUGA) and the D box (CTGA)). Interestingly, even when only fungi closely related to *Cryptococcus* are considered, the sequence of these introns is not well conserved except for the central region containing the snoRNA. For the snoRNAs carried by lncRNAs, this situation is even more remarkable as the only part of the lncRNAs evolutionary conserved is the part bearing the snoRNAs. Actually, the poor exon sequence conservation in these lncRNAs suggests that their roles might be restricted to the production of these snoRNAs. A representative example of a lncRNA-bearing-snoRNA sequence conservation pattern in close *Cryptococcus* relative species is given on the figure 5B. Outside of these close relative species the sequence conservation of the snoRNA sequences is restricted to the consensus boxes.

**Figure 4.**
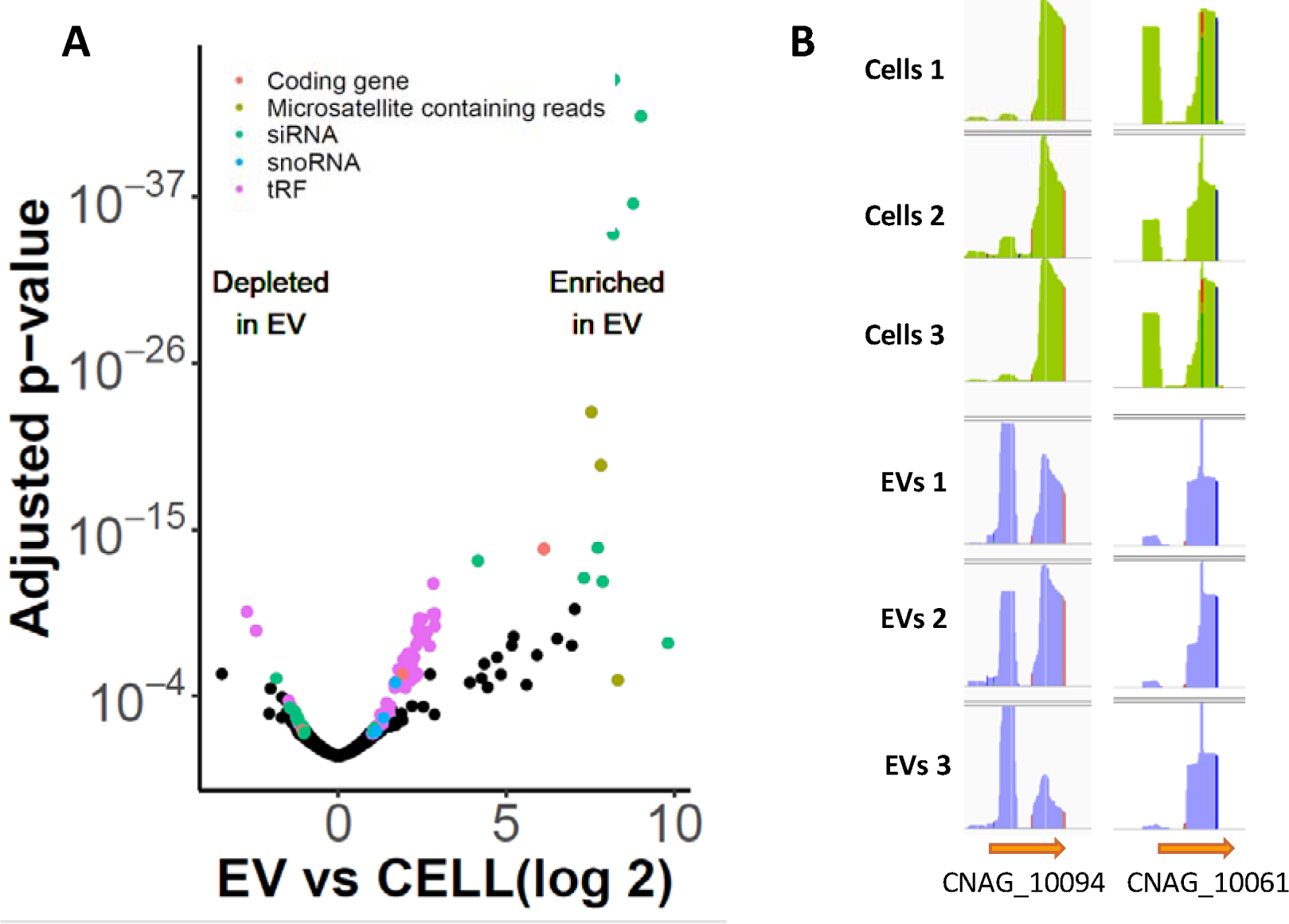
Bias in EV vs cell small RNA content. (A) Volcano plot illustrating the difference between the small RNA transcriptome in EVs and the cells producing these EVs (B) IGV visualization of examples of alteration of the 5’tRF vs 3’tRF repartition in EVs compared to cells at two tRNA loci. Three replicates of the experiment are shown.

**Figure 5.**
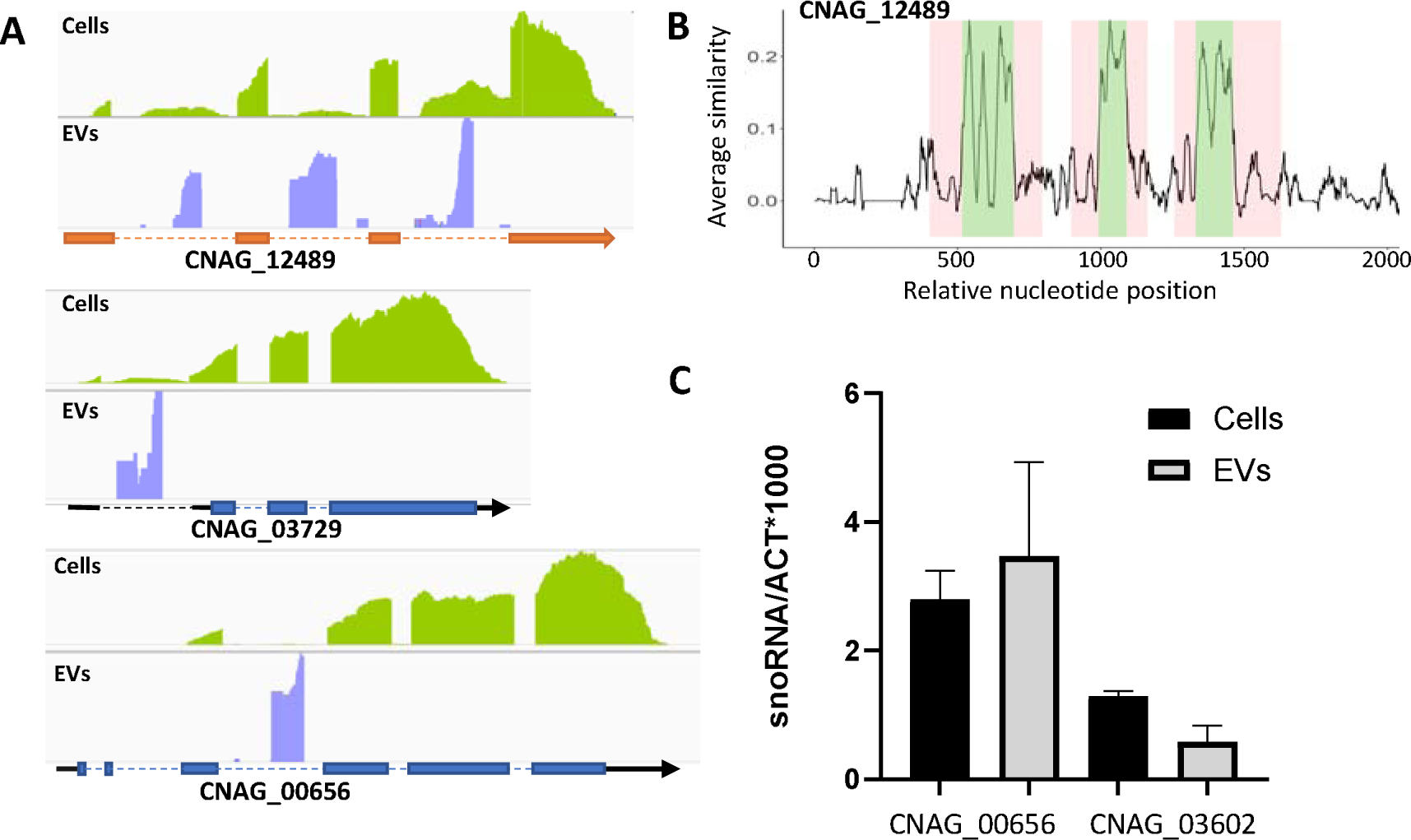
SnoRNAs are associated with EVs in *Cryptococcus*. (A) Examples of snoRNAs associated coverage as observed by IGV. The lncRNA CNAG_12489 encodes three intronic snoRNAs. The coding genes CNAG_03729 and CNAG_00656 contain a snoRNA in a 5’UTR and CDS intron, respectively. (B) Sequence alignment of close *C. neoformans* relative species (*C. deneoformans, C. deuterogattii, Kwoniella heveanensis, Kwoniella bestiolae and Kwoniella dejecticola*) orthologous loci, revealed that the sequence conservation is restricted to the portion corresponding to the snoRNAs within the introns. The green regions correspond to the snoRNA sequences whereas the intronic sequence outside are highlighted in pink (C) snoRNAs do not seem enriched in EVs as revealed by RT-qPCR.. Experiments were conducted in triplicate on RNAs extracted from EVs or cells producing EVs.

We also identified 29 snoRNAs with H/ACA boxes. However, whereas 83.6% of snoRNAs with C/D boxes are in introns, most H/ACA boxes (55.2%, n =16) are located within lncRNAs and one in the intron of a lncRNA. Few of them were located in 5’UTR (n=2), 3’UTR (n=4) or introns (n=4) of coding genes. One snoRNA was overlapping the CDS and the 3’UTR of CNAG_03007 (*CPC1*) suggesting an annotation issue. The remaining snoRNA H/ACA was located within an intergenic region between the CNAG_06094 and the CNAG_06095 loci. We here annotated it as CNAG_13211. In these cases, the sequence conservation with other fungi outside of close relative is also restricted to the consensus boxes. Visual comparison on IGV of our dataset from EVs or EVs-producing cells (Rizzo *et al*. 2023) suggested that snoRNAs might be enriched in the EV RNA samples (Figure 5A).

However, RT-qPCR experiments performed on total RNA purified from EVs and EVs-producing cells did not confirm this impression, highlighting the need of experimental confirmation in this type of analysis (Figure 5C). Analysis of the *C. deneoformans* and *C. deuterogattii* EV sequencing data revealed a similar profile and a very high conservation of these sequences in the three *Cryptococcus* species.

In the EV transcriptome, we also found reads associated with lncRNAs with no annotated function. We manually curated the corresponding annotation and performed RFAM analysis to identify their putative function (Data S1). We identified this way the five main spliceosome components. U1 (CNAG_12993), and U2 (CNAG_12573) were structurally well annotated but had no assigned function. U4 (CNAG_13128) annotation was changed to fit the reads alignment profile. U6 (newly annotated locus CNAG_13208) located in the region between CNAG_05444 and CNAG_05445 was not annotated. U5 was present within the 5’UTR region of the coding gene CNAG_01093, but the alignment profile of the reads from both EVs and cells suggested that the transcription of U5 and CNAG_01093 might be independent. This is supported by the presence of a thymine stretch at the 3’ end of the putative U5 locus, which is a characteristic of RNA polymerase III transcription termination. We thus reannotated this locus, CNAG_13209 being U5. We also identified the loci CNAG_12266 and CNAG_12660 as the RNA moiety of the RNases P and MRP, respectively. Finally, we annotated the 7SL ncRNA at new locus CNAG_13210.

### PolyA RNAs in EVs

We also prepared RNA-seq libraries from polyA+ RNA of EV samples in triplicate. Because the first replicate yielded a very low number of reads, it was subsequently excluded from further analysis. Reads were then aligned to the *C. neoformans* H99 reference genome (Janbon *et al*. 2014) and the number of reads aligned to each annotated feature was counted (Wallace *et al*. 2020). We used this counting to compare the mRNA and lncRNA abundances in EVs and in cells producing these EVs (Rizzo *et al*. 2023). Our analysis did not reveal spectacular enrichment of any mRNA in EVs (Figure 6). Among the genes with at least 100 reads in both EV and cell samples, the most enriched mRNA (5.06-fold) in EVs (CNAG_07756) encodes a *C. neoformans* ubiquitin-protein transferase activating protein Cdc20 homolog. It is important to note that this analysis was limited by the fact that number of genes and lncRNAs have no read in at least one EV or cell replicate. Overall, as no outliers were identified, it is difficult to draw definitive conclusions regarding specific mRNA or lncRNA enrichment in EVs.

**Figure 6.**
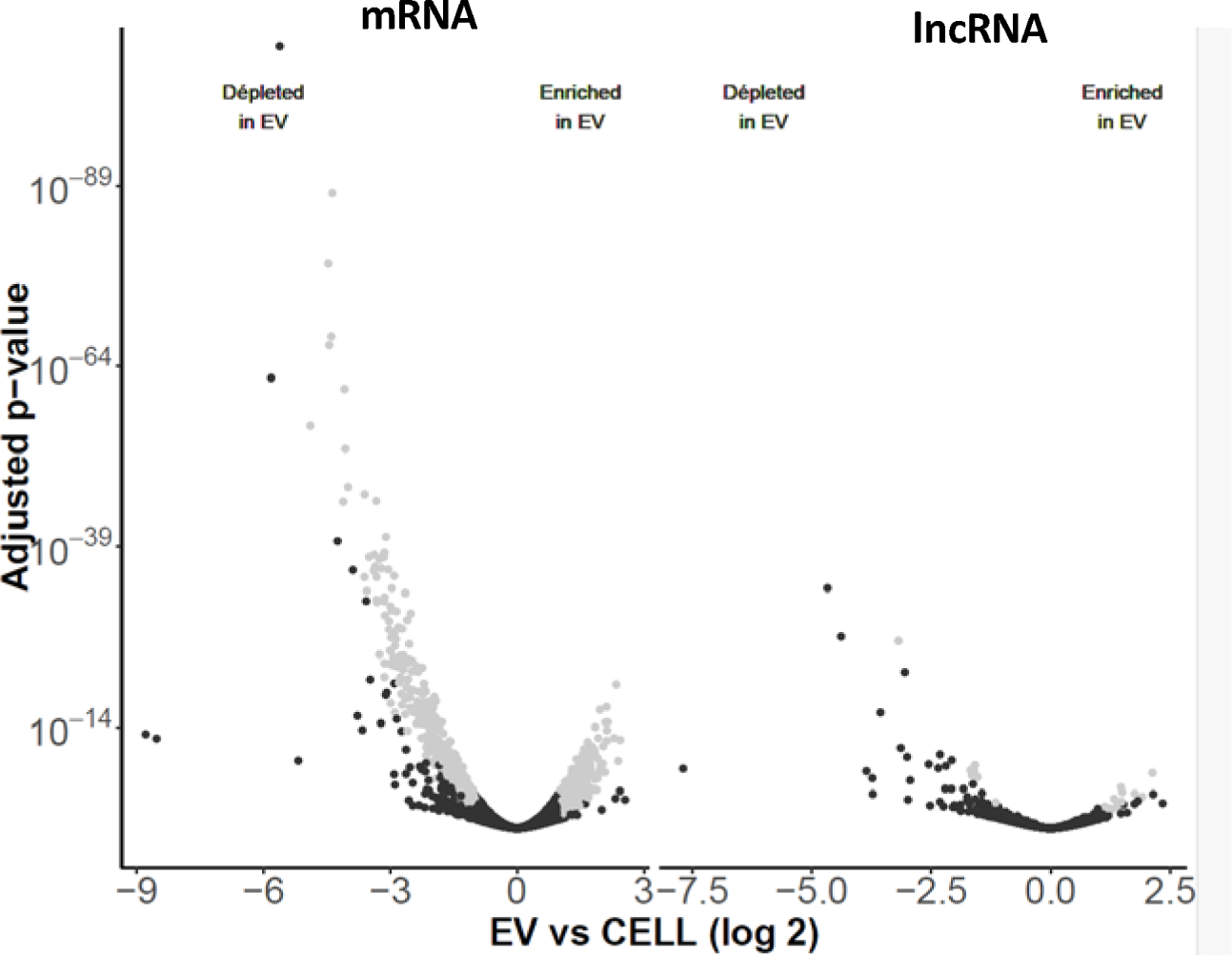
Volcano plot illustrating the difference between the polyA+ RNA transcriptome in EVs and in the cells producing these EVs

## Discussion

We and others have produced a large quantity of data describing the structure and the dynamics of the *Cryptococcus* cellular transcriptome (Janbon *et al*. 2014; Gonzalez-Hilarion *et al*. 2016; Gröhs Ferrareze *et al*. 2021; Kalem and Panepinto 2022). However, even though the study of extracellular transcriptome has become a hot topic in number of organisms including microbes (Hulstaert *et al*. 2020; Pita *et al*. 2020; Gruner and Mcmanus 2021), the diversity of RNA molecules secreted by these yeasts remain poorly described. In fungi, the extracellular transcriptome has been mostly studied through the analysis of RNA molecules associated with EVs (Bitencourt *et al*. 2022b). Several papers reported the presence of small RNAs as well as polyA+ RNAs associated with fungal EV extracts (Bitencourt *et al*. 2022b). However, due to the heterogeneity of the protocols used to prepare these extracts, the diversity of strategies employed for sequencing library preparation and data analysis, as well as the general absence of experimental validation, the overall picture remains blurry. We here used a standardized and validated protocol to prepare EV extracts from three *Cryptococcus* species, we demonstrated that EVs and RNAs co-sediment in an iodixanol based density gradient. We purified and sequenced these RNAs in parallel with RNAs extracted from the very cells producing these EVs. We finally analyzed this data using a structurally and functionally improved genome annotation. None of the data we have produced is entirely new and they mostly strengthened and improved previous results obtained by other teams. As previously reported (Peres Da Silva *et al*. 2015), we identified a large number of tRF reads in the vesicles transcriptome but we here used an improved version of the tRNA annotation in *C. neoformans* to better quantify them. Our analysis also reveals for the first time that at certain tRNA loci, the proportion of 5’tRF vs 3’tRF is altered in EVs compared to cells suggesting specific targeting mechanisms. The origin of the tRFs in *Cryptococcus* is still unknown as no angiogenin (Kumar *et al*. 2016) homolog can be identified in these species. Interestingly, the siRNA deficient strain has conserved a Dicer homolog encoding gene in his genome (D’souza *et al*. 2011) but its role in the biogenesis of the tRFs remains to be studied. Similarly, the presence of reads specific of snoRNAs in fungal EV-associated small RNAs sequencing data was already reported (Peres Da Silva *et al*. 2015). We here extended this information by omitting the size selection step in a small RNA library sequencing kit protocol. Our results allow us to complete the characterization of the *Cryptococus* snoRNAome (Canzler *et al*. 2018; Kalem and Panepinto 2022). Yet, we did not observe a specific enrichment in EV extract for the two snoRNAs we checked by RT-qPCR. It does not mean that these are contaminants and that they have no function in EVs. For instance, the argonaute protein WAGO which has been reported to play a major role in EV biology in the parasitic worm *Heligmosomoides bakeri* is not particularly enriched in EVs compared to the cell (Chow *et al*. 2019). Similar data has been published in mammals in which EV-mediated communication is supposed to rely mostly on miRNA exchange (Weaver and Patton 2020). Besides, the H/CA ribonucleoprotein complex subunit 3, Nop10 (CNAG_01049) was found in our EV proteomic analysis (Rizzo et al 2023) which is in agreement with an active targeting of some snoRNAs to EVs. Our research also confirmed the very poor conservation of snoRNA sequences in fungi. It also suggested that some lncRNA unique function may be to encode snoRNAs in their intron. Finally, we confirmed the presence of siRNAs associated with *C. neoformans* EVs which has been previously reported in both *C. neoformans* and *C. deneoformans* (Peres Da Silva *et al*. 2015; Liu *et al*. 2020). Their structure and the fact that we cannot identify them in the RNAi-deficient species *C. deuterogattii* strongly suggested that these are genuine siRNAs (D’souza *et al*. 2011). As expected, and in agreement with the *Cryptococcus* literature, siRNA reads aligned to transposons or transposon-like sequences (Janbon *et al*. 2010; Dumesic *et al*. 2013). Here also, the analysis of the proteome data revealed that the unique *C. neoformans* Argonaute protein is associated with EVs suggesting an active targeting of siRNAs. The presence of siRNAs associated with EVs is not unique to *C. neoformans* (Peres Da Silva *et al*. 2015) and seems to be a general pattern in fungi. In the recent literature, some of these siRNAs have been identified as miRNAs (Peres Da Silva *et al*. 2015; Liu *et al*. 2020). However, identifying miRNAs in a fungal species by aligning sequences to mammalian miRNAs remains a questionable strategy. In fact, the re-analysis of miRNAs and miRNA-like molecules identified in fungi has demonstrated that none of these newly miRNAs identified passed the various quality criteria (Johnson *et al*. 2022). At this stage, a miRNA-like role of some of these siRNAs cannot be ruled out without experimental confirmation.

The functions of these EV-associated RNA molecules remain unknown, but the literature is full of examples of cell-to-cell communication mechanisms based on RNA molecules exchanges within the same species but also between species including in host-pathogen interactions (Tsatsaronis *et al*. 2018; Cai *et al*. 2021). Although *C. neoformans* EVs have been deeply studied for their structure and composition, their very role in the communication between cells but also during the interaction with the host remain to be uncovered. Examples of EV-mediated exchanges of RNA species (such as siRNAs, miRNAs, tRFs or snoRNAs) between cells are flourishing both in pathogenic and non-pathogenic settings (Weiberg *et al*. 2013; Buck *et al*. 2014; Rimer *et al*. 2018; Ren *et al*. 2019; Sahr *et al*. 2022). These results suggest that this type of exchanges might exist as well in *Cryptococcus* although the characterization of the associated mechanisms remain very challenging.

## Acknowledgements

We here want to thank Eugene Gladyshev and Sebastian Castro Ramirez (Institut Pasteur) for their help in the construction of small RNA libraires. We want Mariette Matondo and Thibault Chase from the Institut Pasteur proteomic facilities for proteomic data production and analysis. The work on EVs in the Janbon’s laboratory is supported by the ANR grants ResistEV (AAPG2021 CE35) and FEVCOM (AAPG2023 CE35). A.T. is funded by a PhD Fellowship from Ecole Doctorale BioSPC, Université de Paris, 75006 Paris, France.

---

Table S1: Proteomic analysis of iodixanol gradient fractions

Table S2: Genes targeted by EV-siRNAs in *C. neoformans*

Table S3: list of the loci enriched in EVs vs cells in small RNA-seq data.

